# Continent-wide effects of urbanization on bird and mammal genetic diversity

**DOI:** 10.1101/733170

**Authors:** C. Schmidt, M. Domaratzki, R.P. Kinnunen, J. Bowman, C.J. Garroway

**Author notes:** Correspondence to: Chloé Schmidt, Biological Sciences Building, University of Manitoba, 50 Sifton Rd, Winnipeg, MB, Canada R3T 2N2; Colin Garroway, Biological Sciences Building, University of Manitoba, 50 Sifton Rd, Winnipeg, MB, Canada R3T 2N2.

## Abstract

Urbanization and associated environmental changes are causing global declines in vertebrate populations. In general, population declines of the magnitudes now detected should lead to reduced effective population sizes for animals living in proximity to humans and disturbed lands. This is cause for concern because effective population sizes set the rate of genetic diversity loss due to genetic drift, the rate of increase in inbreeding, and the efficiency with which selection can act on beneficial alleles. We predicted that the effects of urbanization should decrease effective population size and genetic diversity, and increase population-level genetic differentiation. To test for such patterns, we repurposed and reanalyzed publicly archived genetic data sets for North American birds and mammals. After filtering, we had usable raw genotype data from 85 studies and 41,023 individuals, sampled from 1,008 locations spanning 41 mammal and 25 bird species. We used census-based urban-rural designations, human population density, and the Human Footprint Index as measures of urbanization and habitat disturbance. As predicted, mammals sampled in more disturbed environments had lower effective population sizes and genetic diversity, and were more genetically differentiated from those in more natural environments. There were no consistent relationships detectable for birds. This suggests that, in general, mammal populations living near humans may have less capacity to respond adaptively to further environmental changes, and be more likely to suffer from effects of inbreeding.

## 1. Background

Human activities are among the most prominent and efficient drivers of contemporary evolution (1). In some cases, human-caused evolution in wild populations is well understood and predictable. For instance, we have a well-founded expectation that populations of pests and disease agents will respond adaptively to our attempts at controlling them (1). It is also clear that humans inadvertently alter evolutionary change in wild populations through land use and habitat degradation (2, 3). Whether the indirect effects of human activities on evolutionary change can cause predictable evolutionary outcomes is less well understood. We hypothesized that human land use, by limiting population size and fragmenting habitat, reduces effective population size and genetic diversity in wild populations leading to increased genetic differentiation. To investigate this hypothesis, we repurposed and reanalyzed publicly archived raw nuclear genetic data sets for North American birds and mammals to test for general relationships between urbanization and the genetic diversity of populations.

Urbanization is one of the most pervasive causes of habitat fragmentation and general landscape change. In addition to the ∼700,000 km^2^ occupied by cities (4), nearly 75% of the Earth’s land surface has been modified by humans, primarily in support of city dwellers (5). This human-caused degradation of the planet’s land surface has consistently reduced its capacity to support wildlife (6). As a result, between 1970 and 2014 vertebrate populations have on average declined in size by ∼60% (6). Reductions in population size at this level should decrease genetic diversity by increasing the strength of genetic drift—allele frequency variation due to the random sampling of gametes from one generation to the next. Indeed, there have been general declines in the genetic diversity of populations since the industrial revolution (7). While genetic drift is a neutral evolutionary process that operates independently of the selective value of alleles, it reduces the efficiency of deterministic evolutionary processes like selection by causing allele frequencies to randomly deviate from expected values. When drift is strong relative to selection, random gamete sampling becomes the predominant cause of allele frequency change. In addition, increased drift and inbreeding can eventually lead to reduced mean fitness in small populations. If wildlife populations living in proximity to humans generally experience reductions in population size and connectivity, and thus increased drift, then they may systematically become less genetically diverse than those living in less disturbed environments. By altering a population’s genetic composition in this way, human-caused environmental change could make evolutionary responses to such change less efficient.

The fragmented nature of cities leads to particular expectations about how evolutionary processes will be altered within them based on population genetic theory (2,8,9) (Fig. 1). Assuming a finite population of constant size with individuals that randomly mate, die out, and are completely replaced by their offspring each generation, populations will lose genetic diversity at a rate inversely proportional to population size. In reality, natural populations always deviate from these assumptions. Fortunately, we can substitute the concept of effective population size for census population size and the predictive utility of the theory holds. The effective population size is the size of an idealized population that conforms to the preceding assumptions and produces the same rate of drift as observed in the measured population. We can think of effective population size as a measure of the rate at which genetic drift causes a population to lose genetic diversity. Nearly all violations of these assumptions cause the effective population size to be much lower than the census population size, underscoring that drift plays a more important role in determining genetic diversity and the efficiency of selection than what might be expected from census population size alone. We predicted that reduced census population size and gene flow in cities would lead to smaller effective population sizes, decreased genetic diversity, and increased genetic differentiation in urban populations.

**Figure 1.**
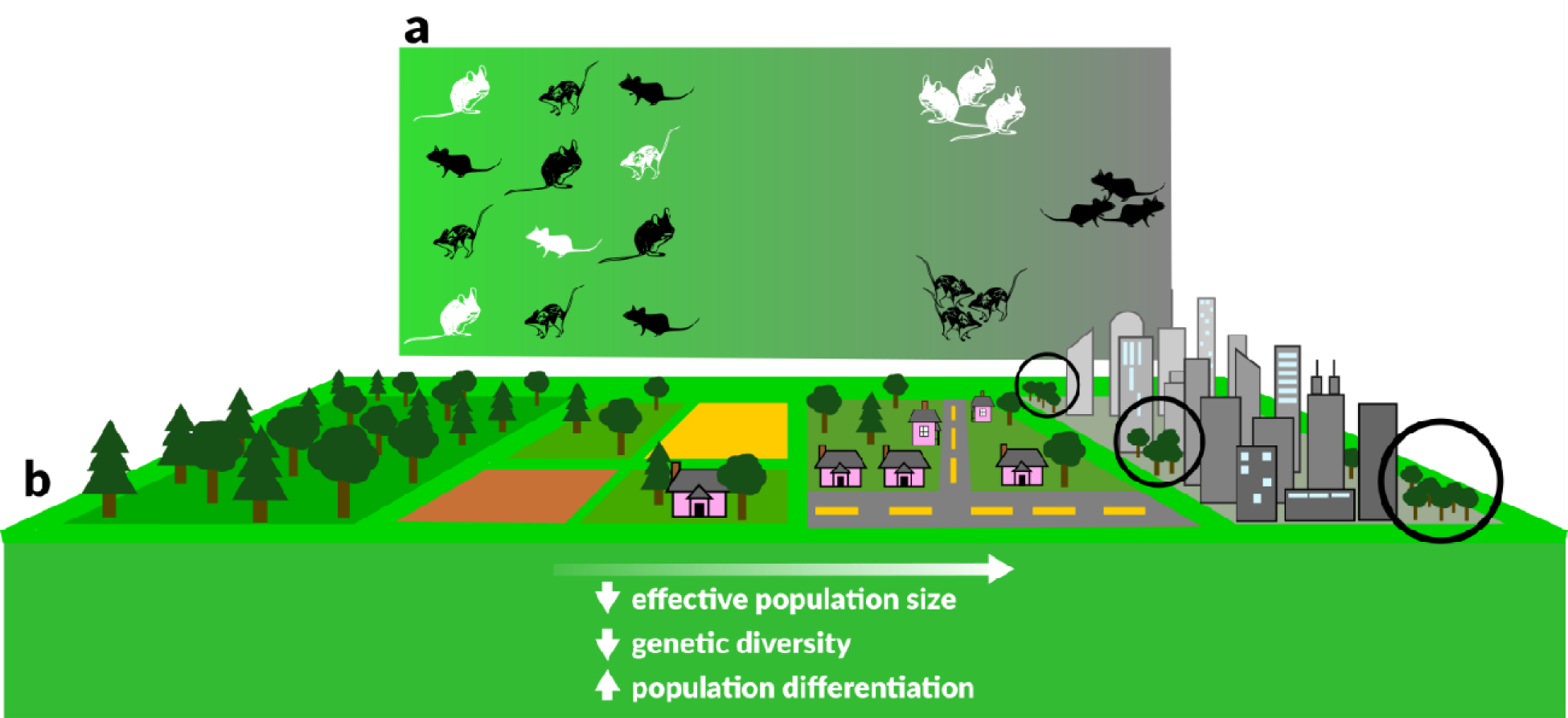
Urbanization is expected to cause smaller effective population sizes, lower genetic diversity, and increased population differentiation in comparison to natural habitats (a). As habitats become increasingly urbanized, they experience greater fragmentation (b), resulting in smaller patch sizes with lower connectivity. Smaller patches limit supportable population sizes wherein genetic drift becomes the predominant evolutionary force and movement between patches in urbanized areas (black circles) becomes difficult, reducing gene flow.

## 2. Collecting and processing raw publicly archived genetic data

We tested for general relationships between the human modification of terrestrial habitats and the genetic composition of North American mammals and birds using archived raw microsatellite data from 85 studies, including 41,023 individuals sampled at 1,008 georeferenced sample sites, spanning 66 species (Table S1, Table S2). In particular, we studied the effects of urbanization and the human footprint (10). We conducted a systematic search of online repositories for all bird and mammal microsatellite data available for North America and applied a series of filtering steps (see SI Methods, Fig. S5) to build a database of georeferenced neutral genetic diversity in wild populations. Our approach was made possible due to the accumulation of data in public data archives, and a still-changing culture of open data in ecological and evolutionary research. Access to georeferenced raw data originally generated for unrelated purposes allowed for a particularly powerful synthetic analysis. This was because we could calculate population genetic parameters and multiple disturbance measures of interest for our question in a consistent manner, whether or not they were presented in the original publications. In addition, the fact that these data were collected to address different questions reduces the likelihood that study system selection—perhaps a tendency to explore evolutionary responses to humans in systems where such responses are expected—biased our findings.

**Table 1.**
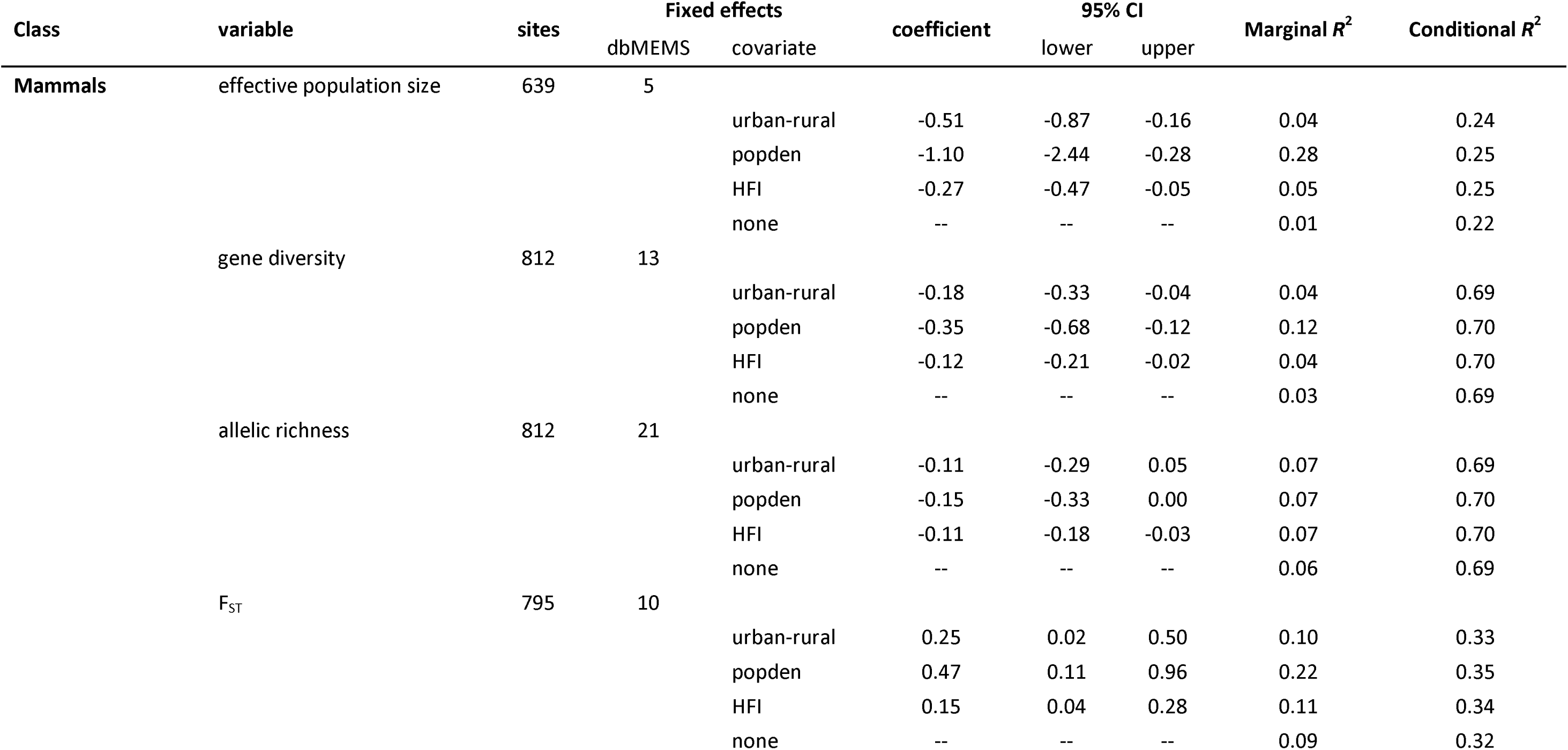
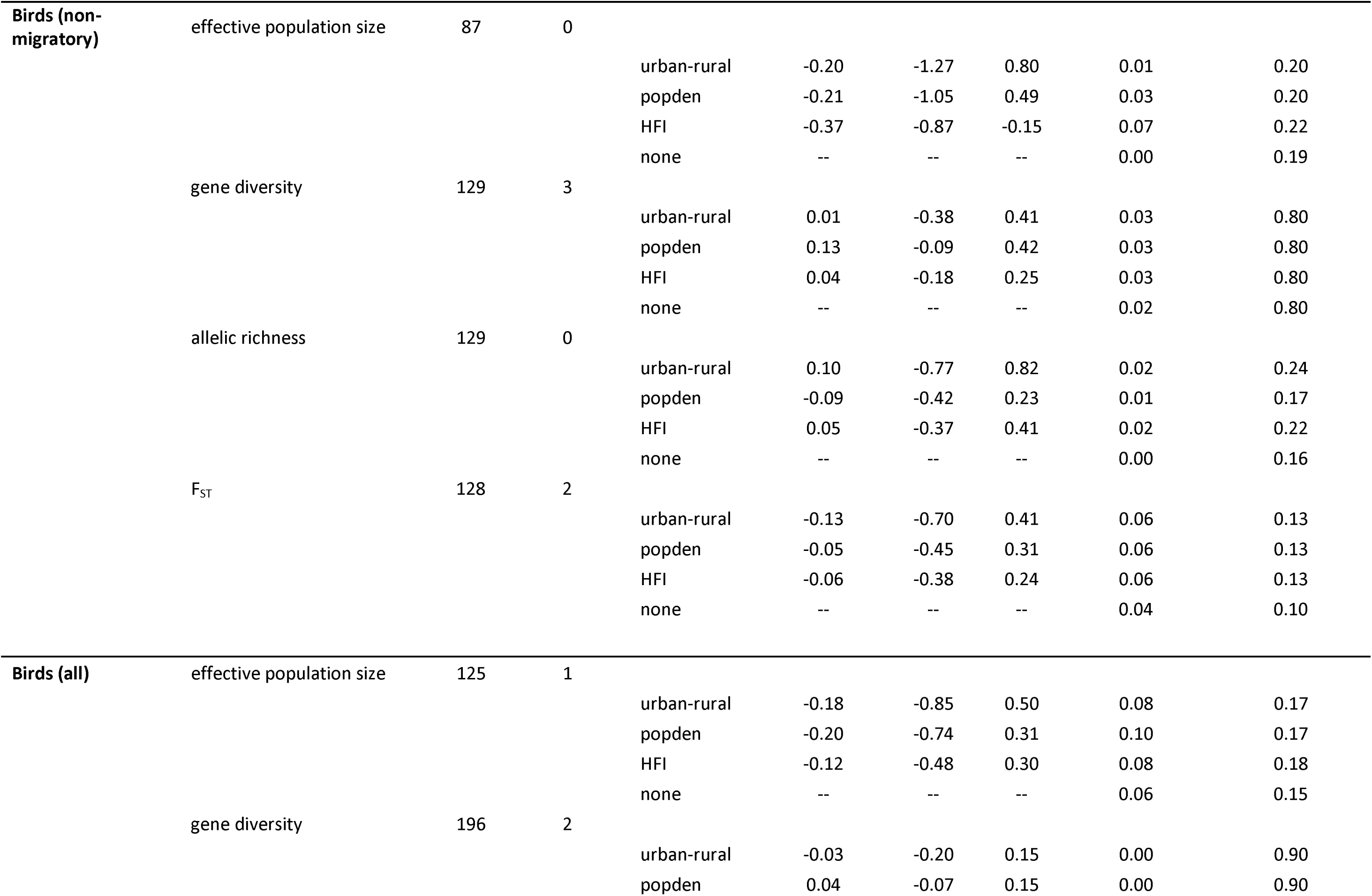
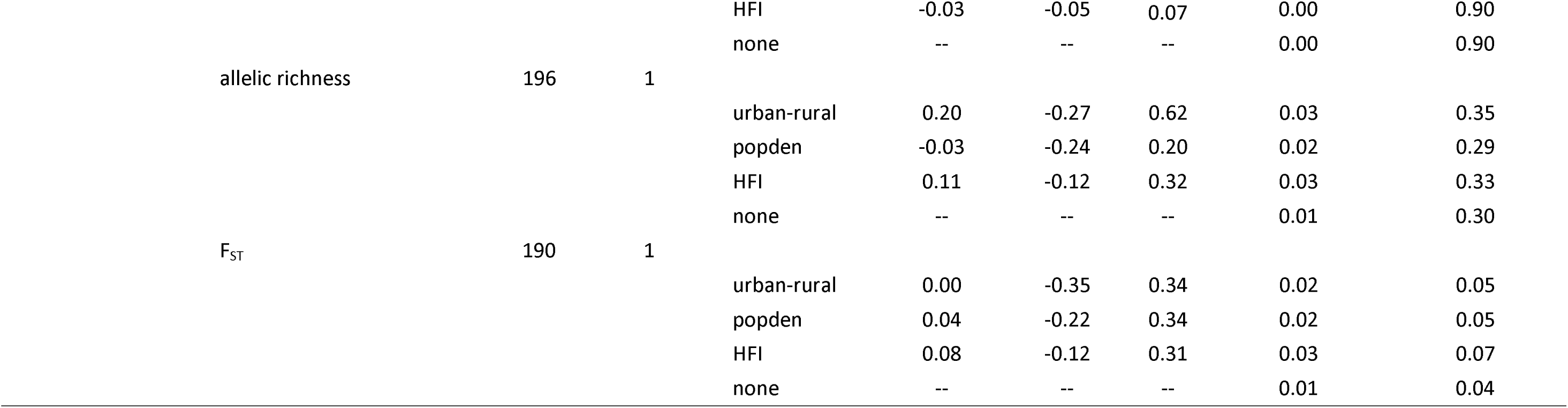
Model summaries for mammals, non-migratory birds, and all birds. Four models were constructed per response variable, each including one of three indices of human disturbance: urban-rural category, human population density (popden), and Human Footprint Index (HFI). The fourth model did not include any measure of disturbance and had only dbMEMs as fixed effects (spatial model), or, where no dbMEMs were selected, a null model. Coefficient of variation, *R*^2^, values are an indicator of model fit; marginal *R*^2^ describes the proportion of variation explained by fixed effects, while conditional *R*^2^ is the variation explained by both fixed and random effects.

We chose to analyze data sets that used neutral microsatellite markers because microsatellites were the most common molecular marker type available in data repositories (11), and because the evolutionary processes that we are interested in are best measured with neutral markers. Although the number of loci surveyed in microsatellite studies is often small relative to surveys of genome-wide markers, the typical number of microsatellites used (∼10 loci) in fact estimates genome-wide diversity well with little gain in accuracy with additional genotyping (12). Variation in microsatellite loci will likely capture recent, fine-scale changes in population structure due to their high mutation rates and variability. While questions about adaptive genetic variation are also interesting, adaptive diversity is currently more difficult to generally define and interpret than neutral genetic diversity, and there are still relatively few data sets suitable for this type of multi-population and multi-species analysis.

We tested for effects of urbanization and the human footprint on estimates of four population genetic parameters calculated for each site: effective population size, gene diversity, allelic richness, and F_ST_ (196 bird sites, of which 129 sampled non-migratory species and were reanalyzed separately, and 812 mammal sites, Fig. 2; Tables 1, S1). We estimated contemporary effective population size of the parental generation using a single sample linkage disequilibrium method to quantify genetic drift (13–15). Of available methods, this approach is one of the more accurate and it is relatively robust to departures from underlying assumptions about population structure (16). Estimators of effective population size perform poorly when sampling error swamps signals of genetic drift, and this meant that effective population size was not estimable at some sites, which we excluded from analysis (17 sites excluded for mammals, 1 site excluded for non-migratory birds; see Table 1 and SI Methods for details). Gene diversity (17) is a measure of genetic diversity that accounts for the evenness and abundance of alleles, and it is not significantly affected by sample size or rare alleles (18). We calculated rarefied allelic richness, the number of alleles per locus corrected for sample size, as a second measure of genetic diversity (19). To quantify genetic differentiation among sites, we estimated site-specific F_ST_ (20). At least two sample sites in a given dataset were required to estimate site-specific F_ST_ (20).

**Figure 2.**
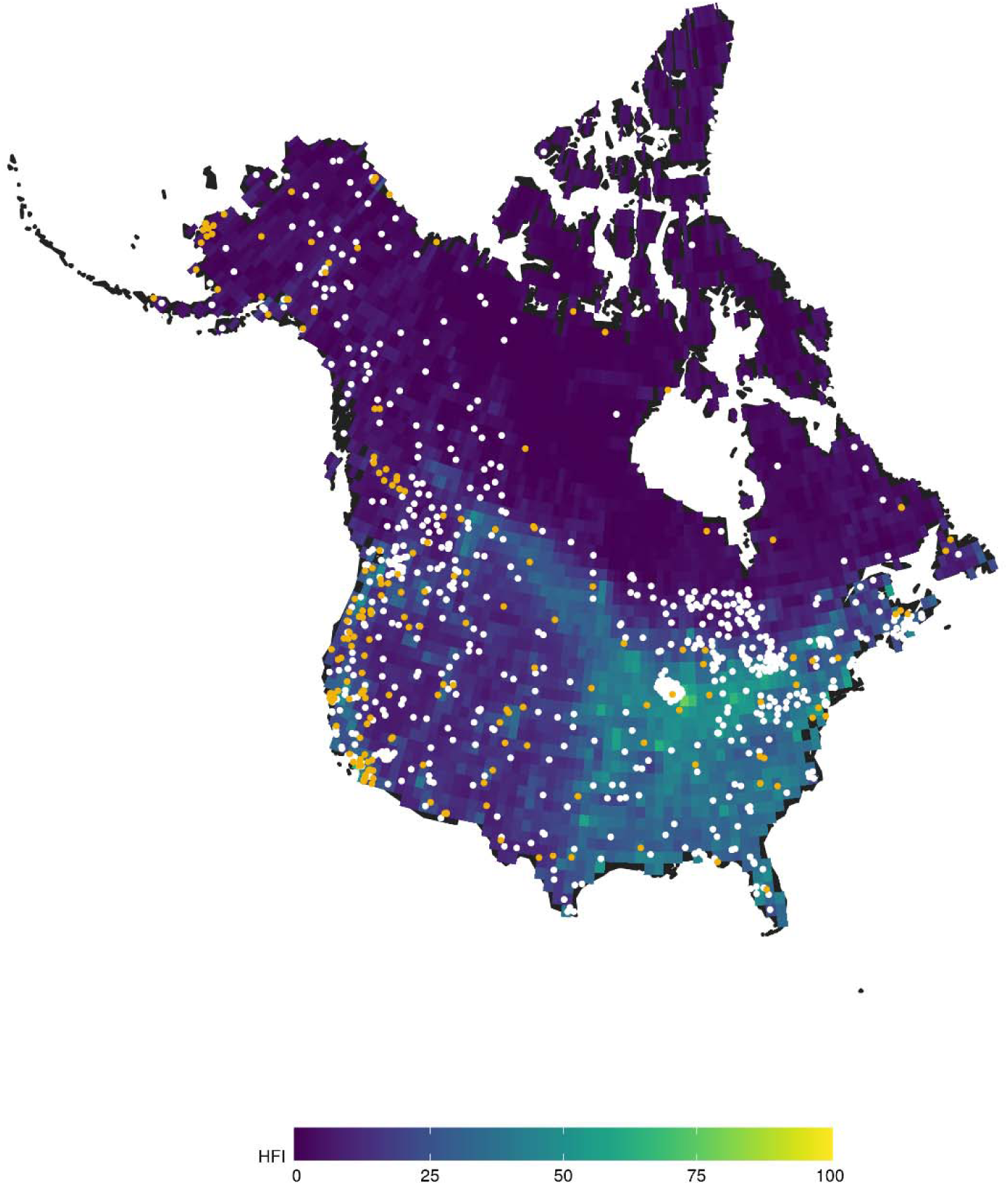
Map of 1,008 sample sites for the 66 mammal and bird species native to North America examined in this study. 812 sites were mammals (white points) and 129 birds (orange points). Using microsatellite markers, we calculated effective population size, gene diversity, allelic richness, and population-specific F_ST_ for each site. Sites are overlaid on a map of the Human Footprint Index (HFI) where values range from 0 (wild habitat) to 100 (disturbed habitat). Note HFI resolution was reduced for the purposes of visualization.

## 3. Modelling strategy

We focused our analyses on the continental United States and Canada due to the historical and demographic similarities of cities and land-usage in this region (21), and to ensure that species have had broadly similar exposures to past climate variation (22). We chose three indices of urbanization and human presence. First, we classified a sample as coming from an urban or rural site based on United States Census Bureau (23) and Statistics Canada (24) classifications of urban areas and population centers. Second, we measured human population density at each site, which may capture aspects of the continuous nature of the effects of human presence that would not be apparent in the binary urban-rural classification. Lastly, we used the Human Footprint Index (10) as a measure of human presence because it incorporates data from multiple land use types including human population density, built-up areas, nighttime lights, land cover, and human access to coastlines, roads, railways, and navigable rivers.

Current levels of genetic diversity will reflect many past processes in addition to urbanization and human-caused environmental degradation. Such processes include exposure to Pleistocene glaciations as well as species-specific life history traits, such as body mass and longevity, each of which could shape effective population size and thus genetic diversity. Because exposure to past environments (22, 25) and life history trait variation (26) vary spatially, we expect the effects of such processes to create spatial variation in genetic diversity. We can account for such spatial patterns by including variables describing spatial patterns in genetic diversity directly in our models, even when the variables themselves are unmeasured (27). This can be accomplished with distance-based Moran’s Eigenvector Maps, or dbMEMs (27–29). Briefly, dbMEMs are orthogonal spatially explicit eigenvectors that summarize spatial autocorrelation (Moran’s I) patterns in data across all scales. In our regression models we used dbMEMs that described spatial variation in our measures of genetic composition to explicitly account for processes causing spatial patterns in the data (29, 30). Neutral genetic diversity also varies with species life history traits which may lack spatial structure (31). We therefore included species as a random effect, allowing both slopes and intercepts to vary, in a generalized mixed modeling framework to capture variation in genetic diversity not already accounted for by dbMEMs (see SI Methods for details).

We used Bayesian generalized linear mixed models to test for relationships between genetic diversity and urbanization (32, 33). We treated each of our four population genetic parameters (effective population size, allelic richness, gene diversity, and site-specific F_ST_) as dependent variables in a series of regression models. Each genetic parameter was fit to each urbanization variable (urban-rural, human population density, and Human Footprint) in separate models that also contained terms for species as a random effect, and spatial variables (dbMEMs) when they were important descriptors of spatial patterns in genetic data. Finally, we fit a null model to each population genetic parameter that contained only the random effect for species and spatial variables. The defining feature of such hierarchical models is that they are models of models – parameter estimates and intercepts can be estimated for each species, and the distribution of these species-specific estimates allows us to generalize effects of urbanization across species. We fit these models for bird and mammal data independently. Migratory behavior in birds may affect spatial patterns in genetic diversity depending on where samples were taken, and whether they were sampled during the breeding season. Therefore, we also ran these models separately for non-migratory birds (7 species, 129 sites; Table S1).

## 4. Results and Discussion

Relationships between all measures of urbanization and the genetic composition of mammal populations were consistently in the predicted directions (see the position of parameter estimates and the breadth of 95% credible intervals in Fig 3a). Effective population size, allelic richness, and gene diversity tended to be negatively related to the measures of urbanization, and sites sampled in areas with greater human presence tended to be the most genetically differentiated (Fig. 3a; Table 1). Contrasting these trends, we found no clear evidence for consistent effects of urbanization and the human footprint on the genetic composition of non-migratory bird samples when analyzed alone (Fig. 3b; Table 1), or when migratory and non-migratory species were combined for analyses (Fig. S1).

**Figure 3.**
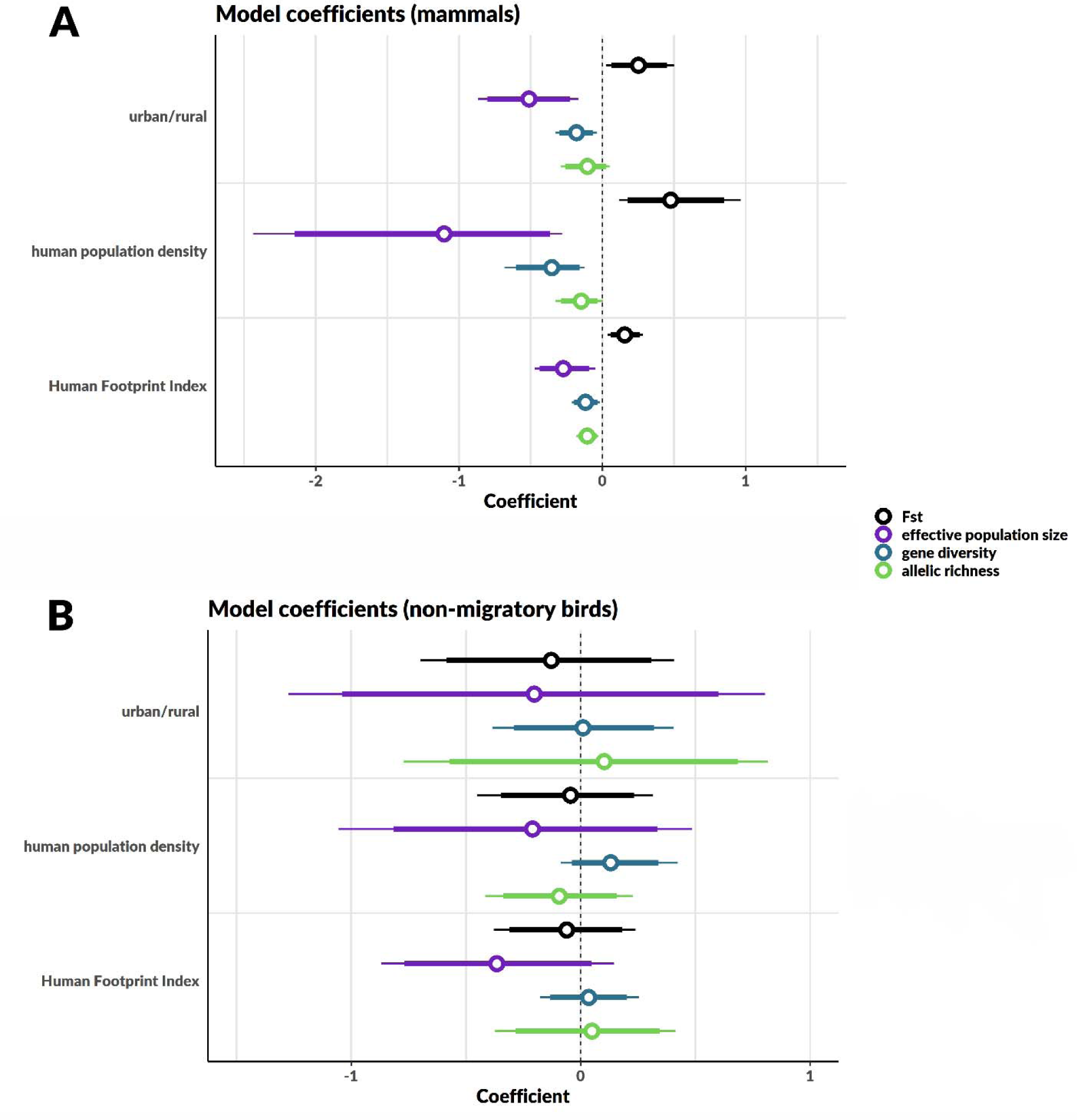
GLMM coefficients for fixed urban and human disturbance effects in mammals (top), and non-migratory birds (bottom; for bird results including migratory species, refer to Table 1 and Fig. S1). Open circles represent coefficient estimates, bold lines are 90% credible intervals, and narrow lines are 95% credible intervals. Sample size differed between variables, i.e. sites where effective population size was not calculable were excluded, and calculation of population-specific F_ST_ for all sites within a study required at least two sample sites. Sample sizes are given in Table 1.

To assess model fits we estimated marginal *R^2^* (*R^2^_m_*), the variance explained by the fixed effects, and conditional *R^2^* (*R^2^_C_*), the variance explained by both fixed and random effects (34). For mammals, all models containing indices of human disturbance explained more variation in the genetic composition of populations than null models (Table 1). Human population density explained the most variation in each measure of the genetic composition of mammal sample sites (effective population size: *R^2^_m_* 0.28; *R^2^_c_* 0.25; gene diversity *R^2^_m_* 0.12; *R^2^_c_*; allelic richness *R*^2^*_m_* 0.07; *R^2^_c_* 0.70; F_ST_ *R^2^_m_* 0.22; *R^2^_c_* 0.35) except allelic richness, where explained variance was similar among all urban predictors (Table 1).

The lack of evidence for genetic effects of urbanization on birds may in part be due to the limited number of data sets available compared to data availability for mammals. Data from seven species remained after excluding migratory species: the California scrub jay (*Aphelocoma californica)*, black-capped chickadee (*Poecile atricapillus)*, boreal chickadee (*P. hudsonicus)*, barn owl (*Tyto alba)*, cactus wren (*Campylorhynchus brunneicapillus)*, spotted owl (*Strix occidentalis)*, and Ridgway’s rail (*Rallus obsoletus).* These species have distinctive ecological and life history traits which vary such that we might not expect to find consistent effects of human presence for these data (Fig. S3). For example, the first four species are human commensals whose population sizes may be expected to increase in proximity to humans. Indeed, exploration of the species-specific effects underlying our mixed model suggests that genetic diversity increases with urbanization for each of these species except barn owls (Fig. S3). In contrast, the Ridgway’s rail and spotted owl are specialized to California salt marshes and old growth forests, respectively, and thus may respond negatively to human presence. However, our results could also be attributable to birds’ vagility. Cities and their surrounding areas are characterized by disjoint patches of habitat interspersed among paved surfaces, buildings, and grassy or agricultural areas (35). Birds’ ability to fly may buffer against the effects of habitat fragmentation and allow for gene flow from undisturbed populations (36) in situations where mammal movements would be more restricted. Indeed, a global analysis of 57 mammal species found that the movements of individuals living in areas with a high Human Footprint Index were considerably reduced relative to those in less disturbed areas (37), which suggests that fragmentation could underlie the patterns we detect in mammals.

In addition to an overall pattern of reduced gene flow and stronger drift in mammals, we note that individual species varied in the strength, and sometimes direction, of their responses to urbanization in our analyses (Fig. S2). This variation could be due to differences in ecological or life history traits which render species amenable to or susceptible to urbanization (11). Surprisingly, we found even synanthropic mammals (e.g. red foxes, *Vulpes vulpes*, and mule deer, *Odocoileus hemionus*) were negatively affected by urbanization at a genetic level (Fig. S2).

The only other studies that have synthetically reanalyzed raw molecular genetic data from online repositories to test for effects of human land-use on genetic diversity have used mitochondrial DNA (mtDNA). Miraldo et al. (38) and Millette et al. (39) reanalyzed raw mtDNA sequence data at a global scale across multiple taxa to assess spatial patterns of variation in sequence diversity, and to explore their relationships with measures of human disturbance. These studies arrived at somewhat contradictory results. Miraldo et al. (38) looked at sequence variation in two mitochondrial genes in mammals, *cytochrome b* and *cytochrome oxidase subunit I*, and found that while genetic variation in *cytochrome oxidase subunit I* increased in less disturbed environments, *cytochrome b* variation was not obviously related to human disturbance. Millette et al.’s more recent work (39) spanned more taxa, including both birds and mammals, and examined variation across spatial scales as well as temporal variation in genetic diversity as measured at *cytochrome oxidase subunit I*. Interestingly, they detected no overarching trend for a loss of genetic diversity associated with proximity to humans (measured by human population density), and no systematic decline in diversity through time. They found considerable spatial, temporal, and taxonomic variation in diversity trends.

How can we explain our results in light of this previous work? Our first thought is that differences could be due to the general lack of relationship between mtDNA diversity and population size (40). Habitat fragmentation and reduced population sizes are hypothesized to be the leading mechanisms causing reduced genetic diversity – if these processes are not captured well by mtDNA markers, trends may be difficult to detect. Additionally, the studies by Miraldo et al. (38) and Millette et al. (39) were global in scope. This is certainly a strength of their work, but underlying spatial differences may be harder to detect and control for at this scale. By focusing on North America, we attempted to control for variation in the timing and nature of disturbances that would otherwise be difficult at a global scale. Taken together, it is clear that there are interesting spatial trends in genetic variation, and the exploration of their underlying causes warrants further study.

While to our knowledge there have not been other synthetic analyses of raw nuclear genetic data similar to ours, there have been syntheses based on published measures of genetic variation (11, 41). DiBattista (41) found that allelic richness and heterozygosity (an amalgam of expected and observed) tended to be lower for mammals described as coming from disturbed populations relative to those from undisturbed populations and, similar to us, found no differences for bird species. Miles et al. (11) also found that allelic richness tended to be reduced in urban populations across a diverse array of taxa. Expected heterozygosity and F_ST_ tended to be lower in urban areas, but this trend was not universal and not significant. They also noted clear instances where urbanization appeared to facilitate gene flow and diversity. We detected some positive effects of urbanization (Figs. S2, S3), but our results more strongly suggest a general negative effect of urbanization on the genetic composition of mammal populations. We suspect that differences in effect sizes and consistency between our work and these previous studies are partly due to our access to raw georeferenced genetic data sets. This meant that we could take a population-level approach, calculating the same genetic measures for all sites, and comparing them to consistently measured indices of disturbance across the entire data set. For example, we could avoid procedures that might add noise to relationships such as combining two measures of heterozygosity and, by calculating a site-specific F_ST_ metric, not have to take means of pairwise measures. An advantage of using literature-based data is that the sample of species was higher than ours for each of the literature-based syntheses. Regardless of approach, these syntheses and ours paint a consistent picture of the effects of urbanization on genetic diversity.

Urbanization and the broader human footprint are leading causes of the current high rates of species and population-level biodiversity losses (42, 43). The population genetic patterns we detected reflect patterns in genome-wide nuclear genetic diversity that are ultimately the result of disturbances related to human presence at the ecosystem, community, and population levels. The consistent effects for mammals across our three measures of human disturbance suggest that this pattern is not confined to just urban spaces – human land use is an issue for the genetic diversity of species in general. This has considerable importance for understanding the current nature and future consequences of biodiversity loss. While monitoring individuals within populations is the best tool for detecting population trends and assessing risk, such direct monitoring of many species is not logistically possible. Our results suggest that calls for genetic monitoring programs are warranted and feasible (44). A relatively small number of genetic markers reflects genome-wide diversity well (12) and is capable of detecting the effects of human presence. In fact, publicly archiving data with publications, originally intended to ensure data posterity, safe-keeping, and research reproducibility, could be better utilized for this task. Access to raw molecular data sets will continue to increase for the foreseeable future, and can be used for monitoring regardless of the original purpose. However, for this to be a useful component of genetic monitoring, more researchers will need to adhere to the standards and best practices for data sharing to maximize reusability (45, 46). This includes using standardized file and metadata formats that are clearly communicated in data package metadata, and including all relevant methodological information. In the data searches presented here, a majority (192/313 = 61%) of datasets were excluded because they did not meet our study criteria (Data S1). However, an additional 36 datasets were excluded for reasons associated with difficulty accessing or interpreting data (Data S1): for example, not being able to download files (e.g., only metadata was available, or only select datasets were deposited), or unclear methodological detail (e.g., no species designations, delineation between study groups was unclear, or lack of spatial reference). We were able to resolve such issues in many cases by contacting the authors, however this might not always be practical for larger studies and limits the ability to automate the data collection process.

Relative to populations in more natural environments, mammal populations in proximity to humans have a reduced capacity to spread beneficial alleles in response to selection pressures, have reduced genetic diversity which can reduce mean population fitness (47, 48), and are more genetically isolated from natural populations. We are extensively and irreversibly creating environmental change while simultaneously reducing the capacity of some populations to evolve in response. Reducing fragmentation and facilitating population connectivity are therefore key to preserving genetic diversity in mammals. Current estimates suggest that by 2050, just 10% of the planet’s surface will be unaltered by humans (6). Land transformation processes are eroding genetic diversity in mammals, compounding direct effects of habitat loss in a way that threatens the long-term existence of populations that persist.

## Acknowledgements

CS, RPK, and CJG were supported by a Natural Sciences and Engineering Research Council of Canada Discovery Grant to CJG. CS and RPK were additionally supported by U. Manitoba Graduate Fellowships, and a U. Manitoba Graduate Enhancement of Tri-council funding grant to CJG. We would like to thank Grace Ji for creating the program we used to format raw data. We also thank Kyle Lefort, Paul O’Brien, Deborah Leigh, the students of the Autumn 2018 section of Evolutionary Biology (BIOL3300) at U. Manitoba, and two anonymous reviewers for their comments on earlier drafts of the manuscript.

## Supplementary Information

### Methods

#### Microsatellite data compilation

Our dataset was comprised of bird and mammal microsatellite data collected from publicly archived, previously published work (Table S2). To create this dataset, we conducted two systematic searches of online databases (Figure S3). We obtained a list of species names for 859 birds and 450 mammals native to Canada and the United States from the IUCN Red List database which includes all species regardless of Red List status. We then queried the Dryad Digital Repository in February 2018 using a python script with the following search terms: species name (e.g. “*Branta canadensis*”), “microsat*”, “single tandem*”, “short tandem*”, and “str”. This search yielded 194 unique data packages associated with papers. A second search was performed in May 2018, this time querying DataOne.org, a network which provides access to data from multiple repositories such as Dryad, the Knowledge Network for Biocomplexity (KNB), and the United States Geographic Survey (USGS). This search was conducted in R using the dataone package (1), a convenient method of querying the DataOne network. Using identical keywords, 237 unique results were generated, 121 of which overlapped with our first search (Figure S3, Data S1).

All data sets were then individually screened for suitability, ensuring: location (Canada and the United States), taxon (native birds and terrestrial mammals), data type (neutral microsatellite markers), and georeferenced sampling (coordinates, maps, or place names). Studies with other factors which may have influenced genetic diversity (e.g. island sites, genetic rescue, translocation, managed or captive populations) were excluded. In total, data from 85 studies were retained for analysis. In a final step, we assured individual sample sites within datasets adhered to our study criteria, and removed those which did not. We maintained the same sample site delineations as in the original work. Criteria for removal from a dataset included island, managed, or captive populations; sites outside of Canada and the United States; and historical samples (where identified). Sites for which we were unable to extract geographic information were also removed, as well as sites with <5 individuals. Any non-neutral microsatellite markers in the data were removed. We note that we compiled all datasets regardless of their location in urban or rural areas, because we determined the level of human disturbance for individual sites using our own criteria which could be consistently applied across all datasets (see “<u>Measures of urbanization and human presence”</u> below). Finally, unique names were assigned to each site, and all datasets were formatted as either STRUCTURE or GENEPOP files and read into R version 3.4.2 (2) using the adegenet package (3).

#### Geographic site locations

Geographic coordinates provided by the authors were used when available (Table S2). Where spatial location was available for each individual sampled, coordinates were averaged. If site names were provided (e.g. “Yellowstone National Park”) with no coordinate reference, we performed a Google Maps search and noted the resulting coordinates. Where applicable, coordinate information was obtained by searching for site names in the Geographic Names Information System (GNIS) or GeoNames database, as was the case for a few datasets from the USGS. In instances where only maps of sampling sites were available, site coordinates were extracted using a reference map in ArcMap version 10.3.1 (ESRI). When georeferencing map images, if sampling locations indicated regions rather than single points, centroid coordinates served as the site location. Centroid coordinates were also calculated as site location for data accompanied by polygon shapefiles as a spatial reference. All coordinates were recorded using the WGS84 (World Geodetic System 1984) coordinate system in decimal degrees, and transformed from other systems or map projections in ArcMap as needed. Finally, when site locations were offshore (42 sites), points were moved to the nearest terrestrial location using the Generate Near Table tool in ArcGIS. Offshore sites (those located in bodies of water) were moved to avoid generating null values for population density and the Human Footprint Index— both of which are high-resolution terrestrial maps which do not extend far past the coast. In some instances, offshore sites were recorded as thus in the original publication, while other times they were generated during the process of obtaining a single location for a site (e.g. the average location of individual coordinates, or centroid location, was in a body of water). Polar bear sites in the Arctic Archipelago constituted half of all offshore sites, while the remainder were coastal species and species sampled near lakes or oceans.

#### Genetic diversity estimates

We chose to measure gene diversity (4) and allelic richness for each site as measures of genetic diversity. Gene diversity uses allele frequencies to determine the probability that pairs of alleles drawn at random from a population are different, and accounts for both the number and evenness of alleles. This measure is minimally affected by sample size and rare alleles (5), and thus is convenient to use when sample sizes are variable, as is the case here. Gene diversity was calculated using the adegenet package (3). Allelic richness, the number of alleles per locus, is strongly influenced by sample size and effective population size. To account for differences in sample size, we used rarefaction as employed in the R package hierfstat (6) to standardize allele counts to the minimum sample size (*n* = 5 individuals) across sites (7). Values were then averaged across loci to obtain a single value per site.

#### Effective population size estimates

We estimated contemporary effective population sizes at each sites using the linkage disequilibrium method for single samples implemented in the software NeEstimator 2.1 (8). The presence of rare alleles produces an upward bias when estimating effective population size which is especially apparent at small sample sizes (9). We therefore set a conservative exclusion threshold (P_crit_) of 0.1, meaning estimates were made based only on alleles with frequencies higher than this value, which has been shown to markedly reduce bias (9). Linkage disequilibrium methods work well for estimating effective population sizes in small populations, however are less reliable for large populations (10). An estimate of infinity is returned when sampling error swamps detectable signals of genetic drift—which may be the case if too few individuals or loci were sampled to yield any useful information about effective population size. In these instances, rather than replacing infinity values with arbitrary large values, we chose to exclude all sites for which we were unable to estimate effective population size (Table 1).

#### Population-specific F_ST_

To estimate levels of population differentiation in relation to human disturbance, we measured population-specific F_ST_ (11). Population-specific F_ST_ characterizes differentiation using the proportion of pairs of matching alleles within populations (the probability of identity by descent) relative to that of pairs from different populations. It can be interpreted as a measure of how far single populations have diverged from a common ancestor population. This measure differs from pairwise F_ST_ estimates (12) in that it provides a measure of population differentiation for a single population, as opposed to a single value for population pairs. Using a population-specific estimator of structure allows us to make comparisons between populations of different species. Population-specific F_ST_ was calculated in R using hierfstat (6), and values were averaged across loci. Because population-specific F_ST_ calculations still use comparisons of pairs of alleles between populations, it could only be measured for species with two or more sample sites. Sample size was slightly decreased when this condition was not met (Table 1).

#### Measures of urbanization and human presence

##### Urban-rural classification

Our next step was to define urban habitats in North America. The United States Census Bureau and Statistics Canada provide publicly available maps of urban areas and population centers, respectively (13, 14). According to the US Census Bureau, an urban area is defined as any densely developed territory with at least 2500 inhabitants. Statistics Canada defines a population center as any area with a minimum population of 1000, and a population density of 400 persons or more per square kilometer. We considered these international designations of urbanization to be comparable. Canadian and American urban area maps were downloaded as polygon GIS layers and merged into a single layer. Site coordinates were transformed from WGS84 to the same projection as the urban area maps (GCS North American 1983) in ArcMap to ensure correct alignment. A spatial join was then performed between sites and the urban area layer in order to classify sample locations as “urban” or “nonurban”. The search radius parameter was set to 10 km to encompass the entire urban gradient, and account for sprawl. Periurban landscapes which are adjacent to cities may be less densely inhabited, however often encompass areas highly managed or disturbed by humans including farmland, parks, and golf courses; in larger cities, periurban landscapes may extend up to 10 km away from the city center (15). Thus, any site located in, or within 10 km of, an urban area was considered “urban” for the purposes of this study.

##### Human population density

Human population density was used as a proxy of urbanization and human effects on the environment. In contrast to our binary urban-rural designation of sample sites, human population reflects the continuous distribution of the effects of human presence, and thus should indicate the intensity of the effects of human activity on genetic diversity. A raster map of global population density per square kilometer was obtained for the most recent available year (2000) from NASA’s Center for Near Earth Object Studies (https://neo.sci.gsfc.nasa.gov/view.php?datasetId=SEDAC_POP). Next, the raster map and shapefile containing sites as point features were read into R (package rgdal and raster; Bivand et al. 2017, Hijmans 2017). Mean population density was calculated within a 10 km buffer zone around each site.

##### Human Footprint Index

The Global Human Footprint Index (18, 19) quantifies human influence on a scale of 0 (most wild) to 100 (most transformed) at a 1 km^2^ resolution. It provides a more comprehensive assessment of the effects of humans than urban-rural designations or population density alone because it incorporates data from multiple sources of land use. In particular, it captures human population density, human land use and infrastructure (built-up areas, nighttime lights, land use, and land cover), and human access (coastlines, roads, railways, and navigable rivers). As with the raster map of population density, the Human Footprint Index was imported to R and values per site (within a 10 km buffer zone) calculated using the same method.

#### Statistical analysis

We modelled birds and mammals separately because we expected them to respond to human disturbance in fundamentally different ways. Within birds, we further classified each species as migratory or non-migratory using information from species accounts in <u>The Birds of North America</u> (20). We then created a separate data subset comprised of only non-migratory species which was analyzed in parallel. Species with a mix of migratory and resident populations were counted as migratory and excluded, as were species with unknown migratory behavior.

Genetic diversity is also affected by regional historical contingencies which would be difficult to specifically identify without detailed knowledge of each species and region in our data set (21). Such events will, however, produce spatial patterns in our genetic measures. These spatial patterns are detectable and can be controlled for—even if their causes are unknown—using distance-based Moran’s Eigenvector Maps (dbMEMs) (22–24). The dbMEM analysis we used (R package adespatial (25)) is a type of eigenanalysis based on principal coordinates analysis which produces a set of spatially explicit variables, dbMEMs, that quantify spatial trends at multiple scales. Because they are orthogonal, dbMEMs can subsequently be included in regression analyses to explicitly model spatial patterns (25). In the first steps of dbMEM analysis, a modified matrix of distances between pairs of sites is calculated from site coordinates. The eigenvalues of this matrix are proportional to Moran’s I coefficients of spatial autocorrelation (23, 26). Importantly, only positive eigenvalues are considered because negative eigenvalues generate complex principal coordinate axes (27). dbMEMs therefore correspond to positive values of Moran’s I, and can account for positive spatial autocorrelation present in the data. Positive spatial autocorrelation occurs when sites nearer to each other are more similar than sites further away, and violates the assumption of independence in our statistical tests. Before undertaking dbMEM, any linear trends in the response variables were removed. Although dbMEM analysis is capable of detecting linear spatial gradients, dbMEMs used to model such trends then cannot be used to recover other, potentially more interesting spatial patterns (22). dbMEM analyses were run in parallel for measures of genetic diversity (gene diversity and allelic richness), population-specific F_ST_, and effective population size. We were able to calculate gene diversity and allelic richness for all sites, however, removed sites where effective population size was infinite and sites where population-specific F_ST_ could not be computed. To capitalize on available data, we created subsets for genetic diversity, population-specific F_ST_, and effective population size, omitting rows where the focal variable(s) had null values. For each taxon we thus had 3 data subsets: one for gene diversity and allelic richness, which included all sites; population-specific F_ST_; and effective population size (Table 1). To select dbMEMs for inclusion in regression analyses, we used forward selection with a p-value criterion (alpha = 0.05) in the SignifReg package (28).

##### Testing effects of human presence on genetic diversity

To test for the effects of human disturbance on genetic diversity, and to determine whether alternate proxies of urbanization would yield similar results, we constructed four linear mixed models per response variable (effective population size, gene diversity, allelic richness, and population-specific F_ST_). Three of these models included spatial dbMEMs and a measure of human presence as explanatory variables: (1) urban-rural category, (2) human population density, and (3) Human Footprint Index. The fourth model consisted of dbMEMs only, or, where no dbMEMs were significant, was a null model (Table 1). Species was included as a random effect in all models to account for species-level variation in genetic diversity, effective population size, and population-specific F_ST_.

The random species effect also accommodated potential variation in the level of species’ responses to human-caused environmental degradation (random slope models). Random effects account for non-independence of samples within groups and increase the accuracy of parameter estimation (29). We fit these models in a Bayesian framework using the R package brms (30) which fits models using Stan. We used default priors (uniform distribution over all real numbers) for parameter estimates with 4000 iterations after discarding warm-up runs (1000 iterations). In cases where models did not converge, we first increased the number of iterations or warmup period (*mammals:* allelic richness ∼ population density: 5000 iterations, 5000 warmup; *birds:* F_ST_ ∼ population density, 5000 iterations, 4000 warmup; *non-migratory birds:* allelic richness ∼ urban category, 4000 iterations, 2000 warmup; allelic richness ∼ population density, 4000 iterations, 4000 warmup; F_ST_ ∼ population density, 4000 iterations, 2000 warmup). If convergence issues persisted we restricted priors to a uniform distribution bounded at −10 and 10 (birds F_ST_ ∼ population density and non-migratory, and birds allelic richness ∼ population density). Lastly, we computed marginal and conditional Bayesian R^2^ to evaluate and compare model fits using the performance package (31).

### Results

#### Data Summary

Our final dataset included 1,008 sites consisting of 66 species (41 mammals and 25 birds; Fig. 2, Table S1). There were 812 mammal sample sites, and 196 bird sample sites (129 non-migratory), with more rural than urban sites (Table S1). The data included samples from a total of 41,023 individuals. Minimum group size for both classes was set to 5 individuals, and there was a maximum of 2444 individuals from a single sample site for mammals (median = 26 individuals), 602 individuals (median = 19 individuals) from a single sample site among all birds, and 141 individuals (median = 19 individuals) for the non-migratory bird subset.

The number of loci sampled ranging between 5 – 210 loci with a median of 13. For all birds (both migratory and non-migratory), the median number of loci sampled was 11 with a range of 6 – 30; non-migratory birds also had a median number of 11 loci with a range between 6 – 20 loci. Gene diversity (mean ± SD) was slightly higher in mammals (0.72 ± 0.11), compared to birds (all: 0.63 ± 0.13; non-migratory: 0.67 ± 0.09). Allelic richness (mean ± SD, max) was similar between mammals (4.79 ± 1.28, 13.70) and birds (all: 4.59 ± 2.19, 22.64; non-migratory: 5.01 ± 2.41, 22.64).

We obtained estimates of effective population size for 639 mammal sites, 125 sites across all birds, and for 87 non-migratory bird sites (Table 1). Effective population sizes (mean ± SD) were on average lower in mammals (614.52 ± 8275.46) compared to birds (all: 980.30 ± 9400.30; non-migratory: 1314.92 ± 11267.02). This corresponded to higher average population-specific F_ST_ among mammals (0.06 ± 0.09) relative to birds (all: 0.04 ± 0.06; non-migratory: 0.04 ± 0.04). F_ST_ was estimated at 796 sites for mammals, 190 for all birds, and 128 for non-migratory birds.

#### Effective population size sample sizes

##### Mammals

We were able to obtain estimates of effective population size for 639 out of 812 mammal sites, with all 41 species represented except for one (moose, *Alces alces*, 2 sites). The ratio of urban to rural sites for sites with non-infinity estimates was unchanged with respect to the full mammal subset (0.44). Additionally, the distribution of sites across values of human population density and the Human Footprint Index did not suggest any bias after removing sites with infinite effective population size estimates (Fig. S4).

##### Non-migratory Birds

Out of 129 sites for non-migratory birds we had 87 non-infinite values. Again, the ratio of urban to rural sites sampled remained consistent with the overall subset of non-migratory birds (0.74), and we saw no indication of bias with regard to human population density or the Human Footprint Index (Fig. S4).

#### Spatial autocorrelation

We found spatial patterns underlying the distribution of genetic diversity in both mammals and birds. dbMEMs capture spatial patterns at all scales in the data, starting broadly (dbMEM 1) and progressing towards increasingly finer scales. In general, we noted more spatial patterns, and more patterns at finer scales in mammals for all response variables. In mammals, following stepwise regression, 5 dbMEMs were significantly related to effective population size (dbMEMs 2, 27, 80, 93, 101). Significant patterns were also found for genetic diversity: 13 dbMEMs were significantly related to gene diversity (dbMEMs 2, 4, 5, 11, 22, 30, 31, 32, 47, 49, 102, 143, 193), and 21 to allelic richness (dbMEMs 2, 4, 5, 7, 8, 11, 12, 13, 21, 22, 29, 30, 31, 32, 47, 49, 102, 108, 143, 185, 190). Finally, we found 10 dbMEMs related to site-specific F_ST_ (dbMEMs 2, 10, 14, 27, 48, 70, 125, 127, 170, 197).

Patterns of spatial variation were less apparent in our subset of non-migratory birds. There were no significant dbMEMs for effective population size nor allelic richness. Gene diversity showed the most spatial variation, with 3 significant dbMEMs selected (dbMEMs 3, 6, 18). For site-specific FST, 2 dbMEMs were significant (dbMEMs 3, 6).

Among all birds, only 1 dbMEMs was significant for effective population size (dbMEM 2), 2 were retained for gene diversity (dbMEMs 2, 6), 1 for allelic richness (dbMEM 2) and1 (dbMEM 6) for site-specific F_ST_.

All significant dbMEMs were incorporated into later models to account for spatial patterns of genetic diversity measures across North America.

---

**Figure S1.**
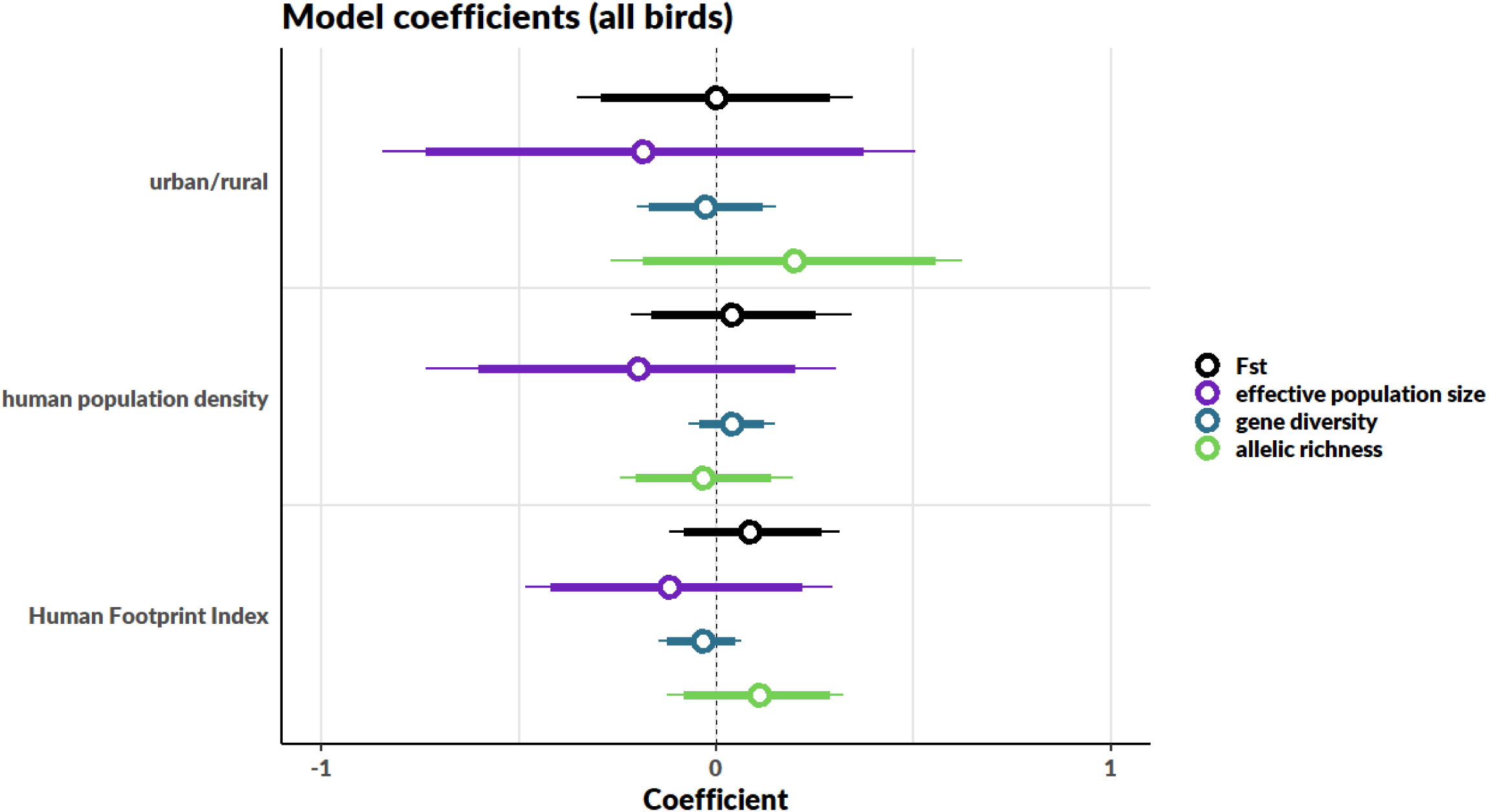
Plotted model coefficients all birds (both migratory and non-migratory species). Open circles represent coefficient estimates, bold lines are 90% credible intervals, and narrow lines are 95% credible intervals. Intervals that overlap zero (dashed vertical line) indicate the disturbance variable has no effect on the response variable. Sample size differed between variables due to limitations estimating effective population size population-specific F_ST_. Sample sizes for each variable are given in Table 1.

**Figure S2.**
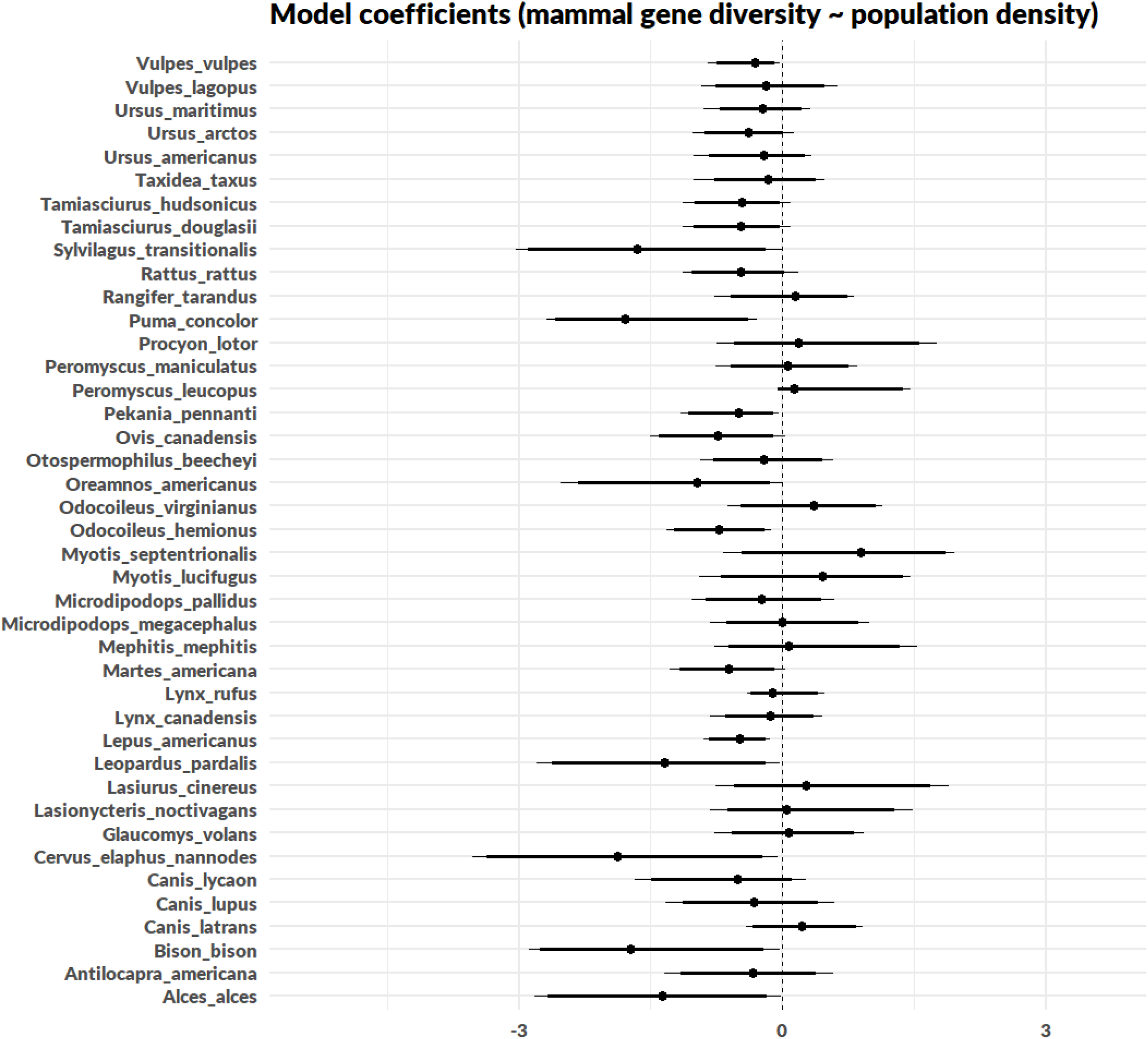
Species-specific parameter estimates for the effect of human population density (which had the strongest effects for mammals) on mammal gene diversity. Ranges are 90% (thick line) and 95% (thin line) credible intervals.

**Figure S3.**
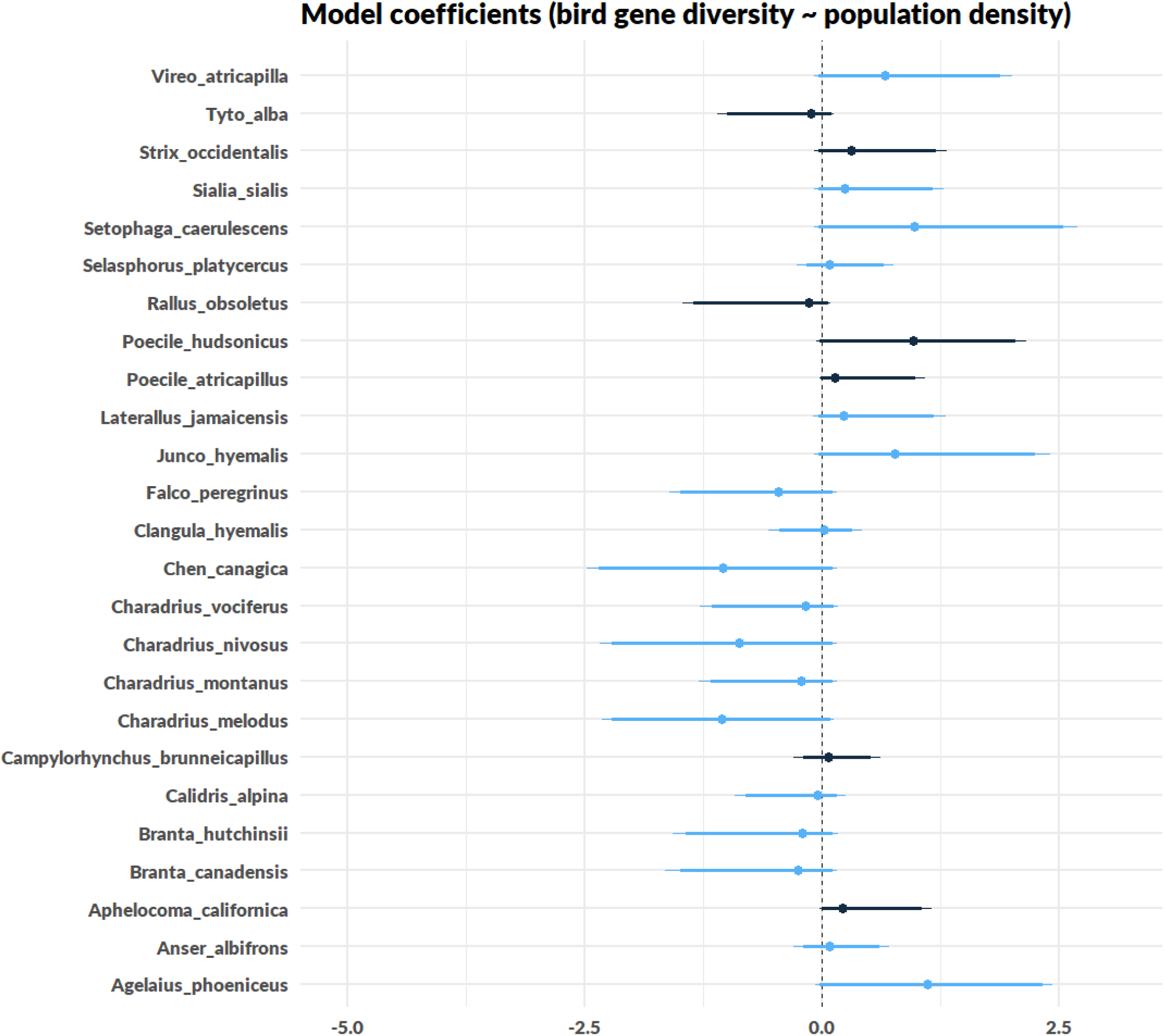
Species-specific parameter estimates for the effect of human population density on gene diversity in migratory (blue) and non-migratory birds (black). Ranges are 90% (thick line) and 95% (thin line) credible intervals.

**Figure S4.**
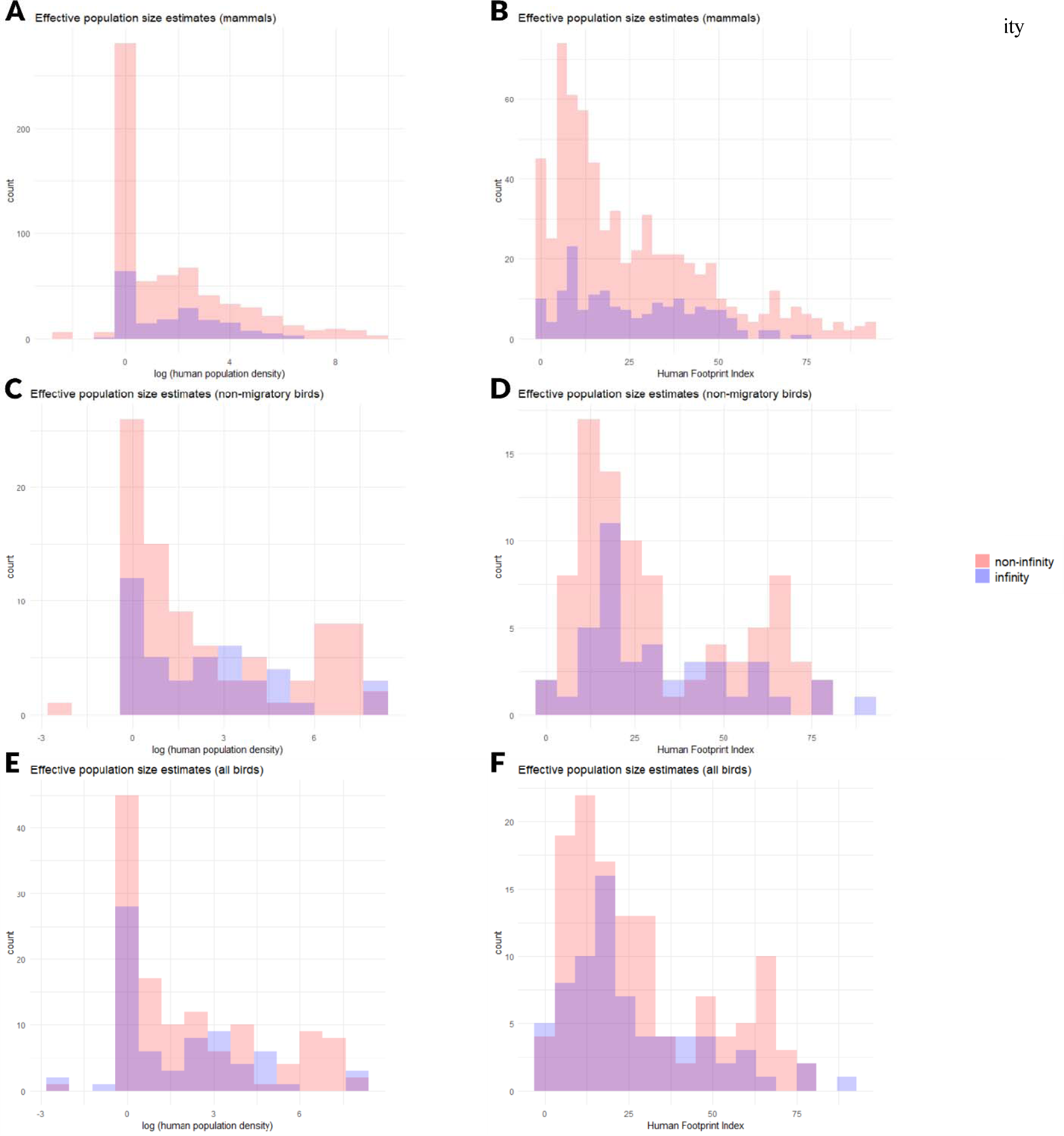
Histograms showing the number of sites included (non-infinity) or excluded (infinity) from effective population size analyses in mammals (top graphs), non-migratory birds (center graphs), and all birds (bottom graphs). (A, C, E) Distribution of sites across human population density. The x axis represents log-transformed human population density plus a constant (0.1). (B, D, F) Distribution of sites across the Human Footprint Index. Human Footprint is measured on a scale from 0 (most wild) to 100 (most disturbed). Overlapping, similar distributions indicate that excluding sites for which we were unable to estimate effective population size likely did not bias our analyses.

**Figure S5.**
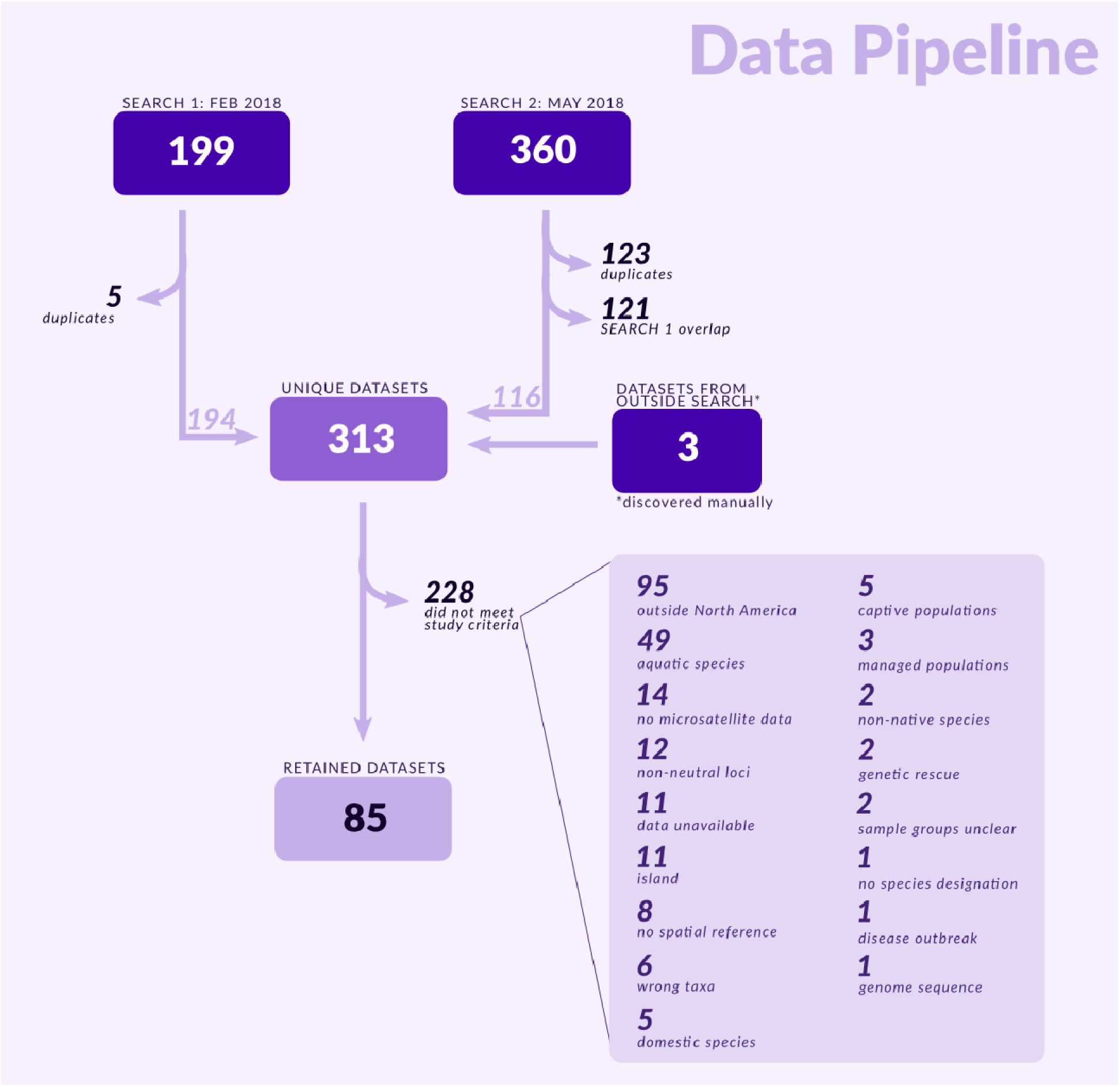
Data pipeline. Schematic shows the number of results from two systematic searches in the DataONE network, and filtering steps taken to arrive at our final dataset. Three microsatellite datasets did not appear in our search results, but were discovered manually and included in our analysis. See Table 2 for a complete list of included work, and Data S1 for raw search results.

**Table S1.** Summary of number of species, sites, individuals and loci per taxonomic class (mammals and birds). For dataset-specific information, refer to Table S3.

| class | species | number of sites |  |  | number of individuals |  | number of loci |  |
| --- | --- | --- | --- | --- | --- | --- | --- | --- |
|  |  | total sites | urban sites | rural sites | median | range | median | range |
| mammals | 41 | 812 | 250 | 562 | 26 | 5 – 2444 | 13 | 5 – 210 |
| birds (all) | 25 | 196 | 71 | 125 | 19 | 5 – 602 | 11 | 6 – 30 |
| birds (non-migratory) | 7 | 129 | 55 | 74 | 19 | 5 – 141 | 11 | 6 – 20 |

**Table S2.** Summary and reference information for data used in this study. **Class**: mammals (M) or birds (B). **Repository**: location of data (USGS = U.S. Geological Survey). **Loci**: number of loci sampled, **populations**: number of populations, **individuals**: total number of individuals summed over all populations. **Coordinates**: method of assigning coordinate locations to each site. C = site coordinates provided in study; AC = coordinates given per sample in study, averaged to obtain site coordinates; GM = site name searched in Google Maps; GNIS = site name searchable in the Geographic Names Information System; M = map provided in study, georeferenced in ArcMap; P = site polygons (shapefile) provided, centroid coordinates taken.

| Species | Class | Repository | Loci | Populations | Individuals | Coordinates | Search Date | Reference |
| --- | --- | --- | --- | --- | --- | --- | --- | --- |
| *Alces alces* | M | Dryad | 10 | 2 | 89 | AC | February | (1, 2) |
| *Canis latrans* | M | Dryad | 10 | 41 | 482 | GM | February | (3, 4) |
| *Cervus elaphus nannodes* | M | Dryad | 20 | 2 | 54 | GM | February | (5, 6) |
| *Lasionycteris noctivagans* | M | Dryad | 18 | 1 | 87 | C | February | (7, 8) |
| *Lasiurus cinereus* | M | Dryad | 19 | 1 | 132 | C |  |  |
| *Lynx rufus* | M | Dryad | 10 | 5 | 95 | GM | February | (9, 10) |
| *Leopardus pardalis* | M | Dryad | 10 | 2 | 70 | GM |  |  |
| *Lynx rufus* | M | Dryad | 9 | 4 | 365 | AC | February | (11, 12) |
| *Lynx rufus* | M | Dryad | 15 | 52 | 1646 | AC | February | (13, 14) |
| *Mephitis mephitis* | M | Dryad | 9 | 1 | 345 | AC | February | (15, 16) |
| *Procyon lotor* | M | Dryad | 10 | 1 | 330 | AC |  |  |
| *Microdipodops megacephalus* | M | Dryad | 11 | 3 | 184 | GM | February | (17, 18) |
| *Microdipodops pallidus* | M | Dryad | 10 | 2 | 105 | GM |  |  |
| *Myotis lucifugus* | M | Dryad | 9 | 15 | 735 | C | February | (19, 20) |
| *Myotis lucifugus* | M | Dryad | 11 | 21 | 1142 | M | February | (21, 22) |
| *Myotis lucifugus* | M | Dryad | 8 | 29 | 1310 | C | February | (23, 24) |
| *Myotis septentrionalis* | M | Dryad | 5 | 15 | 896 | C |  |  |
| *Odocoileus hemionus* | M | Dryad | 10 | 60 | 1831 | C | February | (25, 26) |
| *Oreamnos americanus* | M | Dryad | 22 | 1 | 102 | AC | February | (27, 28) |
| *Ovis canadensis* | M | Dryad | 208 | 1 | 276 | C | February | (29, 30) |
| *Ovis canadensis* | M | Dryad | 210 | 1 | 216 | GM |  |  |
| *Peromyscus leucopus* | M | Dryad | 10 | 12 | 134 | C | February | (31, 32) |
| *Peromyscus leucopus* | M | Dryad | 11 | 11 | 367 | C | February | (33, 34) |
| *Peromyscus leucopus* | M | Dryad | 18 | 13 | 312 | GM | February | (35, 36) |
| *Peromyscus maniculatus* | M | Dryad | 10 | 1 | 31 | GM | February | (37, 38) |
| *Puma concolor* | M | Dryad | 13 | 1 | 739 | AC | February | (39, 40) |
| *Puma concolor* | M | Dryad | 18 | 2 | 667 | AC | February | (41, 42) |
| *Puma concolor* | M | Dryad | 46 | 8 | 354 | M | February | (43, 44) |
| *Puma concolor* | M | Dryad | 10 | 2 | 196 | M | February | (45, 46) |
| *Rangifer tarandus* | M | Dryad | 19 | 18 | 655 | M | February | (47, 48) |
| *Rangifer tarandus* | M | Dryad | 18 | 27 | 606 | M | February | (49, 50) |
| *Rangifer tarandus* | M | Dryad | 21 | 5 | 230 | M | February | (51, 52) |
| *Rangifer tarandus* | M | Dryad | 14 | 27 | 802 | M | February | (53, 54) |
| *Rattus rattus* | M | Dryad | 9 | 8 | 126 | GM | February | (55, 56) |
| *Taxidea taxus* | M | Dryad | 12 | 1 | 233 | AC | February | (57, 58) |
| *Taxidea taxus* | M | Dryad | 12 | 3 | 917 | AC | February | (59, 60) |
| *Taxidea taxus* | M | Dryad | 20 | 8 | 236 | M | February | (61, 62) |
| *Ursus americanus* | M | Dryad | 15 | 7 | 269 | M | February | (63, 64) |
| *Ursus americanus* | M | Dryad | 15 | 4 | 250 | AC | February | (65, 66) |
| *Ursus maritimus* | M | Dryad | 21 | 14 | 2232 | AC | February | (67, 68) |
| *Ursus arctos* | M | Dryad | 8 | 16 | 831 | M | February | (69, 70) |
| *Ursus maritimus* | M | Dryad | 8 | 11 | 319 | M |  |  |
| *Ursus americanus* | M | Dryad | 8 | 1 | 32 | M |  |  |
| *Ursus arctos* | M | Dryad | 20 | 1 | 729 | GM | February | (71, 72) |
| *Ursus maritimus* | M | Dryad | 9 | 2 | 610 | AC | February | (73, 74) |
| *Ursus maritimus* | M | Dryad | 8 | 4 | 402 | GM | February | (75, 76) |
| *Vulpes vulpes* | M | Dryad | 8 | 5 | 257 | M | February | (77, 78) |
| *Vulpes lagopus* | M | Dryad | 9 | 3 | 78 | M |  |  |
| *Vulpes vulpes* | M | Dryad | 13 | 11 | 376 | AC | February | (79, 80) |
| *Canis lupus* | M | Dryad | 12 | 1 | 62 | C | February | (81, 82) |
| *Canis lycaon* | M | Dryad | 12 | 1 | 62 | GM |  |  |
| *Odocoileus_hemionus* | M | Dryad | 10 | 2 | 410 | AC | February | (83, 84) |
| *Tamiasciurus hudsonicus* | M | Dryad | 9 | 12 | 199 | AC | February | (85, 86) |
| *Tamiasciurus douglasii* | M | Dryad | 9 | 14 | 198 | AC |  |  |
| *Lepus americanus* | M | Dryad | 8 | 39 | 853 | AC | February | (87, 88) |
| *Martes americana* | M | Dryad | 12 | 34 | 653 | C | February | (89, 90) |
| *Agelaius phoeniceus* | B | Dryad | 10 | 29 | 240 | C | February | (91, 92) |
| *Aphelocoma californica* | B | Dryad | 13 | 7 | 493 | AC | February | (93, 94) |
| *Calidris alpina* | B | Dryad | 8 | 30 | 236 | C | February | (95, 96) |
| *Charadrius melodus* | B | Dryad | 8 | 3 | 220 | C | February | (97, 98) |
| *Charadrius montanus* | B | Dryad | 14 | 7 | 94 | C |  |  |
| *Charadrius nivosus* | B | Dryad | 15 | 4 | 68 | C |  |  |
| *Charadrius vociferus* | B | Dryad | 14 | 2 | 49 | C |  |  |
| *Laterallus jamaicensis* | B | Dryad | 15 | 2 | 336 | M | February | (99, 100) |
| *Selasphorus platycercus* | B | Dryad | 8 | 2 | 58 | C | February | (101, 102) |
| *Poecile hudsonicus* | B | Dryad | 6 | 2 | 260 | C | February | (103, 104) |
| *Poecile atricapillus* | B | Dryad | 11 | 13 | 913 | C | February | (105, 106) |
| *Poecile atricapillus* | B | Dryad | 14 | 32 | 142 | C | February | (107, 108) |
| *Setophaga caerulescens* | B | Dryad | 6 | 8 | 401 | C | February | (109, 110) |
| *Sialis sialis* | B | Dryad | 12 | 1 | 73 | C | February | (111, 112) |
| *Antilocapra americana* | M | Dryad | 19 | 4 | 175 | GM | May | (113, 114) |
| *Bison bison* | M | Dryad | 29 | 1 | 188 | GM | May | (115, 116) |
| *Odocoileus virginianus* | M | Dryad | 14 | 8 | 2069 | C | May | (117, 118) |
| *Otospermophilus beecheyi* | M | Dryad | 11 | 64 | 205 | AC | May | (119, 120) |
| *Ovis canadensis nelsoni* | M | Dryad | 16 | 3 | 579 | P | May | (121, 122) |
| *Rangifer tarandus* | M | Dryad | 16 | 14 | 480 | AC | May | (123, 124) |
| *Sylvilagus transitionalis* | M | Dryad | 10 | 5 | 157 | AC | May | (125, 126) |
| *Anser albifrons* | B | USGS | 8 | 3 | 115 | C | May | (127) |
| *Campylorhynchus brunneicapillus* | B | Dryad | 22 | 4 | 363 | AC | May | (128, 129) |
| *Clangula hyemalis* | B | USGS | 12 | 12 | 109 | AC | May | (130) |
| *Falco peregrinus* | B | USGS | 12 | 7 | 112 | GNIS | May | (131) |
| *Junco hyemalis* | B | Dryad | 7 | 8 | 602 | GM | May | (132, 133) |
| *Strix occidentalis* | B | USGS | 10 | 1 | 423 | AC | May | (134) |
| *Rallus obsoletus* | B | USGS | 9 | 17 | 107 | C | May | (135) |
| *Vireo atricapilla* | B | Dryad | 12 | 7 | 160 | AC | May | (136, 137) |
| *Ursus americanus* | M | Dryad | 12 | 1 | 2444 | AC | May | (138, 139) |
| *Lynx canadensis* | M | Dryad | 15 | 2 | 556 | M | May | (140, 141) |
| *Lynx rufus* | M | Dryad | 17 | 1 | 106 | AC | May | (142, 143) |
| *Odocoileus hemionus* | M | Dryad | 18 | 5 | 135 | M | May | (144, 145) |
| *Ursus americanus* | M | Dryad | 20 | 4 | 101 | M, AC | May | (146, 147) |
| *Ursus arctos* | M | Dryad | 20 | 4 | 113 | M, AC |  |  |
| *Lynx canadensis* | M | Dryad | 14 | 2 | 702 | C | February | (148, 149) |
| *Glaucomys volans* | M | Dryad | 7 | 2 | 278 | C |  |  |
| *Odocoileus hemionus* | M | USGS | 15 | 28 | 73 | AC | May | (150) |
| *Anser albifrons* | B | USGS | 19 | 8 | 10 | GM | May | (151) |
| *Branta canadensis* | B | USGS | 19 | 1 | 10 | GM |  |  |
| *Branta hutchinsii* | B | USGS | 18 | 1 | 9 | GM |  |  |
| *Chen canagica* | B | USGS | 30 | 1 | 27 | GM |  |  |
| *Ursus americanus* | M | Dryad | 15 | 1 | 506 | AC | May | (152, 153) |
| *Ursus maritimus* | M | Dryad | 24 | 3 | 78 | M | May | (154, 155) |
| *Peromyscus maniculatus* | M | Dryad | 11 | 10 | 109 | C | May | (156, 157) |
| *Pekania pennanti* | M | Dryad | 16 | 28 | 722 | C | NA | (158, 159) |
| *Tyto alba* | B | NA | 20 | 4 | 292 | M | NA | (160) |
| *Vireo atricapilla* | B | Dryad | 9 | 9 | 67 | M | NA | (161, 162) |

## Data S1.xlsx

List of all results from systematic searches performed in February and May 2018. Columns in this table are: class (mammal or bird), species name, search date, repository, whether it was included in our study (yes or no), the reason for exclusion if applicable, and a link to the online data. The “overlap” reason for exclusion indicates May results which overlapped with February results, while “duplicate data” indicates a duplicate entry for the same dataset within a search month. See Figure S5 for our data filtering process. Ultimately, 85 data sets were retained for analysis after individual screening to ensure the data met our study criteria, including: location (within the continental United States and Canada), taxon (native birds and terrestrial mammals), data type (neutral microsatellite markers), and georeferenced sampling. Available at: https://doi.org/10.5061/dryad.cz8w9gj0c

## Notes

#### Summary of Updates

Final submitted draft with minor updates to text.

